# Molecular epidemiology of carbapenem-resistant hypervirulent *Klebsiella pneumoniae*: Risk factors and resistance mechanism of ceftazidime/avibactam in China

**DOI:** 10.1101/2024.11.27.625653

**Authors:** Na Wang, Minghua Zhan, Lexiu Deng, Huiying Li, Xiaocui Peng, Jianliang Chang, Jiatong Hao, Na Jia, Baoliang Li, Chunmei Lei, Teng Wang, Shuting Liu, Caiqing Li, Bu Wang, Wei Zhang

## Abstract

The high morbidity and mortality rates associated with carbapenem-resistant hypervirulent *Klebsiella pneumoniae* have become a global public health problem. We analyzed the whole genome sequence and epidemiological data of 81 carbapenem-resistant *Klebsiella pneumoniae* strains and conducted sensitivity experiments to explore the optimal concentration of carbapenem-resistant *Klebsiella pneumoniae*. We verified the resistance and virulence genes of carbapenem-resistant *Klebsiella pneumoniae* using polymerase chain reaction and compared virulence between carbapenem-resistant hypervirulent *Klebsiella pneumoniae* a ndcarbapenem-resistant-non-hypervirulent *Klebsiella pneumoniae* through *Galleria mellonella* assays.The prevalent types of multilocus sequence typing in our carbapenem-resistant *Klebsiella pneumoniae* strains were ST11 (69.14%, 56/81). ST11-K64-O1/O2v1 has become an epidemic type in our hospital, with drug-resistance genes dominated by bla_KPC-2_ (90.12%, 73/81). Ceftazidime/avibactam has better antibacterial effects on carbapenem-resistant *Klebsiella pneumoniae*, and its resistance is mainly mediated by a resistance plasmid containing the bla_KPC-2_ gene. 52 carbapenem-resistant hypervirulent *Klebsiella pneumoniae* strains carry the bla_KPC-2_, with a high detection rate of siderophore. Further research has shown that the genetic environment of the bla_KPC-2_ contains TnpR_Tn3 and ISKpn27 upstream and ISKpn6 insertion sequences downstream. *Galleria mellonella* revealed that the survival rate of carbapenem-resistant hypervirulent *Klebsiella pneumoniae* was lower than that of carbapenem-resistant-non-hypervirulent *Klebsiella pneumoniae*, and the survival rate of carbapenem-resistant hypervirulent *Klebsiella pneumoniae*-K64 was lower than that of carbapenem-resistant hypervirulent *Klebsiella pneumoniae*-K47.The spread of carbapenem-resistant hypervirulent *Klebsiella pneumoniae* strains is alarming, necessitating molecular monitoring of hypervirulent strains and strengthening the nosocomial infection prevention and control measures. Antibiotics should be used reasonably based on patients’ infection status and local epidemiological data.

**Importance:** Carbapenem-resistant *Klebsiella pneumonia* is a global public health concern. We conducted a molecular epidemiological analysis of 81 carbapenem-resistant hypervirulent *Klebsiella pneumoniae* strains from our hospital and explored the resistance mechanism of ceftazidime/avibactam. We identified the bla_KPC-2_ gene to be primarily responsible for carbapenem resistance in our hospital, and ST11-K64 was the dominant clone. A high-risk clone, ST307-K102, was detected, necessitating strengthened surveillance to prevent its spread. This study’s findings contribute to the optimization of antimicrobial management and indicate that genomic research can assist with the tracking of hospital infection outbreaks and the improvement of infection-control measures.

## Introduction

Carbapenem-resistant *Klebsiella pneumoniae* (CRKP) has become a major global public health threat, and the prevalence of carbapenem-resistant hypervirulent *Klebsiella pneumoniae* (CR-hvKP) has sharply increased in China, leading to high morbidity and mortality rates due to the lack of effective treatment options (1, 2). *Klebsiella pneumoniae* Carbapenemase (KPC) belongs to class A serine carbapenemases and is the main mechanism of resistance in Enterobacteriaceae. The proportion of bla_KPC-2_ producing CRKP in China exceeds 70%(2). In China, ST11 CRKP has become the major clonal type (3), whereas in the United States and Europe, ST258/512 is predominant(4). Owing to ceftazidime/avibactam having good *in vitro* sensitivity and safety against serine carbapenemases, it has become the last line of defense against CRKP infections. However, CRKP can generate bla_KPC_ mutations via various mechanisms, causing resistance to ceftazidime/avibactam(5, 6). Research has found that the diversity of the CRKP genome is mainly due to horizontal transfer, including plasmids, phages, integration, and binding elements. The spread of bla_KPC-2_ is typically mediated by two mobile elements, Tn4401 and NTEKPC (7).

Globally, CR-hvKP is increasingly reported, and its emergence is devastating as CR-hvKP possesses high resistance, virulence, and transmissibility. Most clinical cases occur in Asia, particularly in China, where studies have shown that the prevalence of CR-hvKP has increased from 28.2% in 2016 to 45.7% in 2020 (3), causing serious nosocomial infection in the intensive care unit ward (8). CR-hvKP virulence factors include capsules, siderophores, virulence plasmids, and other virulence genes, which help differentiate hypervirulent strains (9). Various complex mechanisms have led to the rapid spread of strains in the clinic (9). Previous studies have mainly focused on resistant or virulent strains of hvKP, with little attention paid to the correlations among virulence genes, resistance genes, and antimicrobial susceptibility of CR-hvKP. Therefore, this study aimed to summarize the molecular epidemiological characteristics of CR-hvKP isolated from this region and discuss the evolution of virulence and resistance genes in CR-hvKP and their relationship with clinical phenotypes.

## Materials and methods

### 1. Isolate and antimicrobial susceptibility testing

In this study, 81 non-duplicate CRKP strains were collected from the First Affiliated Hospital of Hebei North University between 2020 and 2021 for analysis. The PubMLST database was downloaded for comparative analysis of *Klebsiella pneumoniae* containing bla_KPC-2_ and/or bla_NDM-1_ in November 2024 (https://bigsdb.pasteur.fr/klebsiella/). This study employed the microbroth dilution method recommended by the Clinical and Laboratory Standards Institute to perform *in vitro* antimicrobial susceptibility testing of 81 CRKP strains. Sensitivity, intermediate resistance, and resistance were determined according to the Clinical and Laboratory Standards Institute-M100 ED33 guidelines. A minimum inhibitory concentration (MIC) of ≥4 mg/L for imipenem or meropenem against *KP* was defined as CRKP.

### 2. Polymerase chain reaction

Polymerase chain reaction experiments were conducted on the 81 collected CRKP strains using the following reaction system: The total reaction volume was 25 μL, which included 12.5, 8.5, 1, 1, and 2 μL of 2x Taq Plus Master Mix II (Dye Plus), deionized water, forward primer, reverse primer, and template DNA, respectively. The reaction program was set as follows: initial denaturation at 95°C for 3 min, denaturation at 95°C for 15 s, annealing at 60°C for 20 s, and extension at 72 ° C for 60 s, with a total of 30 cycles. After the reaction, agarose gel electrophoresis was performed to record the experimental results.

### 3. Whole genome sequencing and annotation

Genomic DNA was extracted using the sodium dodecyl sulfate method, and the harvested DNA was analyzed through agarose gel electrophoresis. Quantitative analysis of the DNA was performed using a Qubit® 2.0 fluorometer (Thermo Scientific). Sequencing libraries were prepared using the NEBNext Ultra DNA Library Prep Kit (Illumina, USA), and index codes were added to classify the sequences for each sample. Whole-genome sequencing was performed on an Illumina NovaSeq PE150 platform using a 2 × 150 bp paired-end strategy.

The genome was annotated using Prokka V1.14.6, and resistance, virulence genes, and plasmid replicons were predicted using Abricate V1.0.1. Capsular serotypes were predicted using PathogenWatch (https://pathogen.watch). Multilocus sequence typing (MLST) of all strains was conducted using the Pasteur database (https://bigsdb.pasteur.fr/klebsiella/), and newly identified sequence typing (ST) was submitted to the MLST database administrator for approval and were assigned ST numbers. A minimum spanning tree based on the allelic difference between isolates of the seven housekeeping genes was constructed using PHYLOViZ (10).

### 4. Phylogenetic analysis

Single nucleotide polymorphisms were extracted using Snippy v4.6.0 (https://github.com/tseemann/snippy) to generate a core genome alignment. This core genome alignment was employed to construct a maximum likelihood phylogenetic tree using FastTree V2.1.11-2, with *KP* subsp. *pneumoniae* HS11286 (GCA_000240185.2) as the reference genome. The resulting phylogenetic tree was visualized using iTOL (https://itol.embl.de/) (11).

### 5. Galleria mellonella killing assay

Bacterial suspensions were prepared at concentrations of 10⁴ cfu/mL, 10⁵ cfu/mL, and 10⁶ cfu/mL, and 10 microliters of each were injected into the larvae of the greater wax moth. The control group was injected with an equal volume of phosphate-buffered saline. This procedure was used to establish an infection model for the larvae. To ensure consistency in the infection model, the injection was made approximately 5 mm deep into the second-to-last right legs of the larvae, and the entire volume of the bacterial suspension was injected simultaneously. The infection models were incubated at 37°C, and the number of dead larvae was recorded every 2 h over an observation period of 48 h. Subsequently, the mortality rate of the larvae was calculated.

### 6. Antibiotic-resistant plasmids and mobile genetic elements containing bla_KPC-2_ gene

The assembled contigs of six CR-hvKP strains resistant to cefotaxime/avibactam were input into the VRprofile2 pipeline (https://tool2-mml.sjtu.edu.cn/VRprofile/home.php) [12] to divide the contigs into chromosome or plasmid fragments and identify whether they carry resistance genes. To further identify plasmids carrying resistance genes, we compared these contigs with the NCBI database (https://www.ncbi.nlm.nih.gov/) using blstn to search for reference sequences. For the plasmid resistant to cefotaxime/avibactam, we used pC76 KPC (NZ_CP080299.1) with a coverage of 93.82% and an identity of 99.37% as a reference sequence to study the structure of the plasmid. MAUVE (12) was used to identify all contigs located in drug-resistant plasmids by comparing the contigs of the strain with the reference plasmid pC76 KPC (NZ_CP080299.1). Plasmid maps were presented using BRIG (13).To determine whether contigs carrying the bla_KPC-2_ gene all contain insertion and repeat sequences, we submitted them to Isfinder (14)(https://www-is.biotoul.fr/index.php). Easyfig (15) is used to visualize the structure of the bla_KPC-2_ gene.

#### Statistical methods

In this study, the chi-square test was used to compare differences between the CR-hvKP and CR-non-hvKP groups. The Log-rank (Mantel-Cox) test was used to analyze survival differences between the two groups. The Wilcoxon rank-sum and Kruskal–Wallis tests were used to analyze drug susceptibility differences among the three groups. Statistical significance was set at p-value <0.05. Visualization was performed using Xiantao Academic (https://www.xiantaozi.com/). Spearman’s correlation analysis was conducted to explore the relationships between plasmids and virulence genes, plasmids and resistance genes, resistance genes and clinical data, and virulence genes and clinical data. Heatmap visualization was performed using the Microbiome Cloud Platform (https://www.bioincloud.tech/task-meta). Statistical analyses were performed using IBM SPSS Statistics for Windows, version 27.0 (IBM Corp., Armonk, N.Y., USA), and graphing was performed using Origin 2021 (OriginLab Co. MA, USA) and GraphPad Prism 8 (GraphPad Software, LaJolla, CA, USA).

## Results

### 1. Molecular epidemiology

CRKP has become popular worldwide, with China, the United States, and Brazil being the majority (Figure 2A). In China, CRKP is most common in Beijing, followed by Shanghai and Hunan (Figure 2B). The MLST of CRKP in our hospital is mainly ST11, followed by ST15, which is consistent with the popular trend in China (Figure 3). Our hospital has detected 12 types of ST, among which ST11 (69.14%) and ST15 (11.11%) are the main prevalent types. Capsular (K) antigen serotyping revealed 16 different types, with K64 being the most common (60.49%, 49/81). Lipopolysaccharide (O) antigen serotyping showed that the most common serotypes were O1/O2v1 (66.67%, 54/81). Overall, 81 CRKP strains were primarily isolated from sputum samples (65.43%, 53/81), followed by urine (16.05%, 13/81) and secretion (4.94%, 4/81) samples. Phylogenetic analysis revealed that evolutionary relationships were associated with ST classification and independent of specimen type. Strains of the same ST clustered mostly on the same evolutionary branch, whereas ST11 and ST15 clustered on different branches. The main clusters were ST11-K64-O1/O2v1, ST11-K47-OL101, and ST15-K19-O1/O2v2. The ST11 strains were distinctly categorized into the following three branches: ST11-K64-O1/O2v1, ST11-K47-OL101, and ST11-K25-other O serotypes, suggesting that ST11 strains with various serotypes evolved in different directions (Figures 4 and 5).

**Figure 1.**
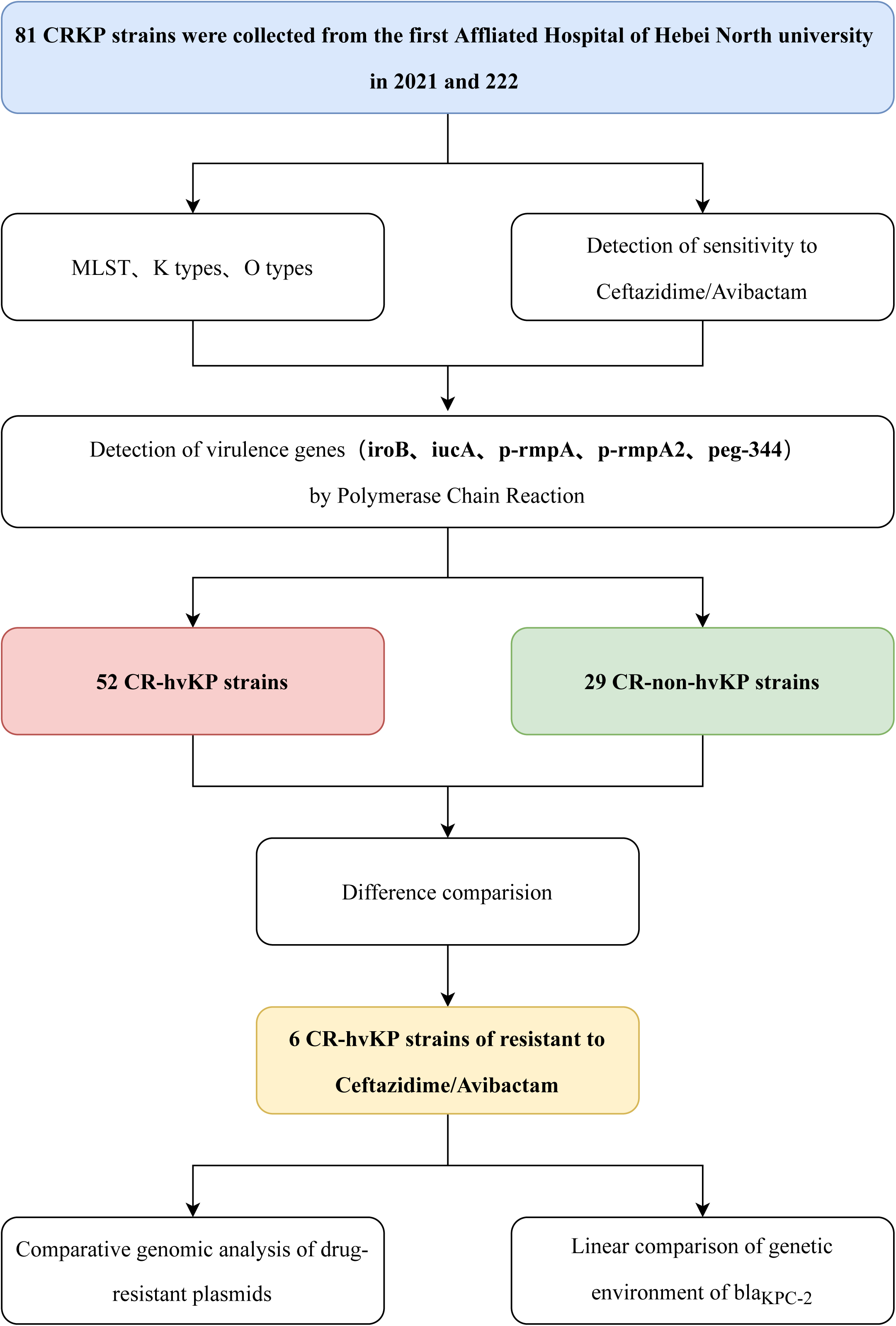
Design and experimental flowchart of this study.

**Figure 2.**
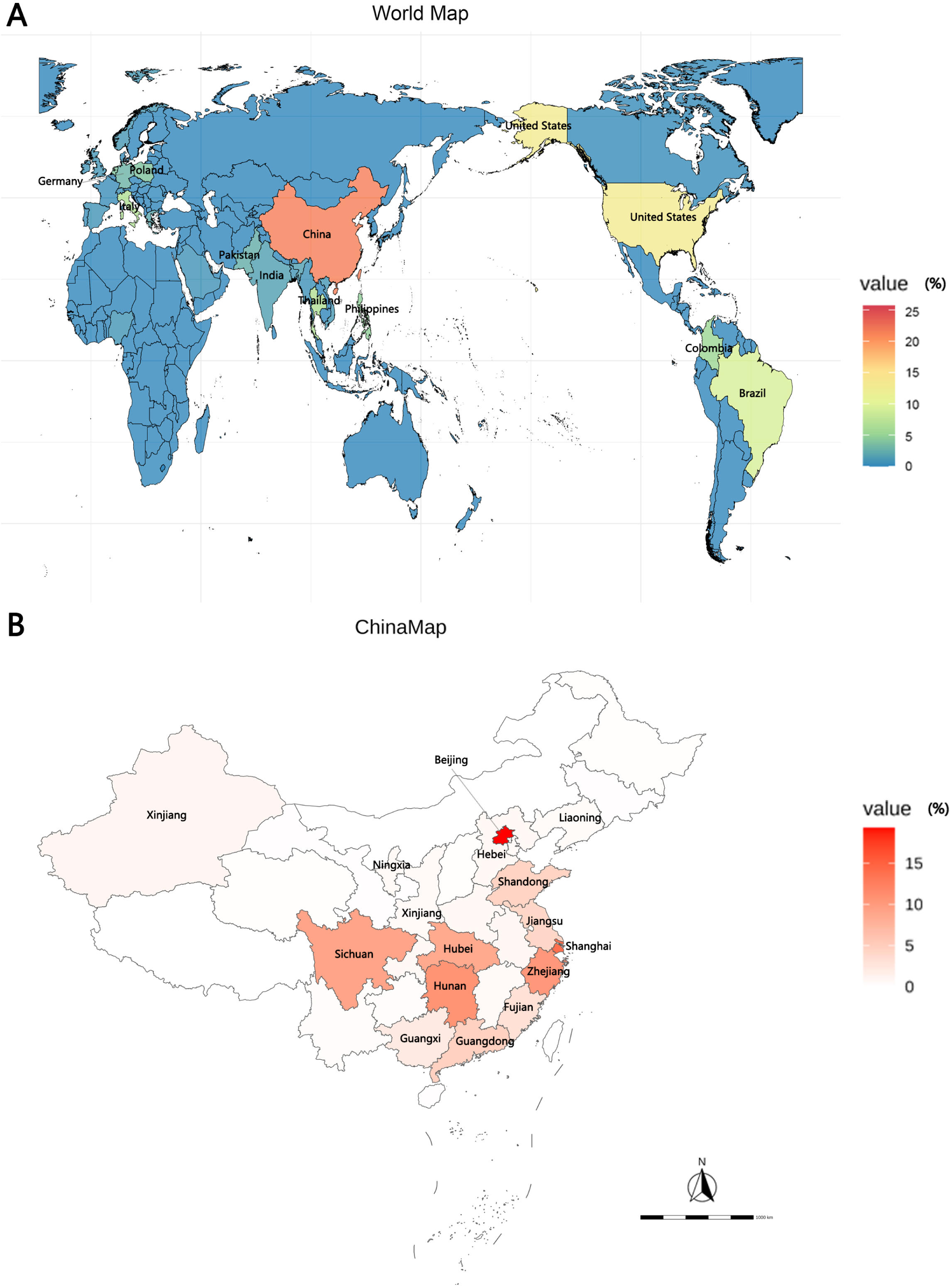
Global and Chinese distribution of CRKP infection by country. A. Global distribution of CRKP infection. B. Chinese distribution of CRKP infection. CRKP, carbapenem-resistant *Klebsiella pneumonia*

**Figure 3.**
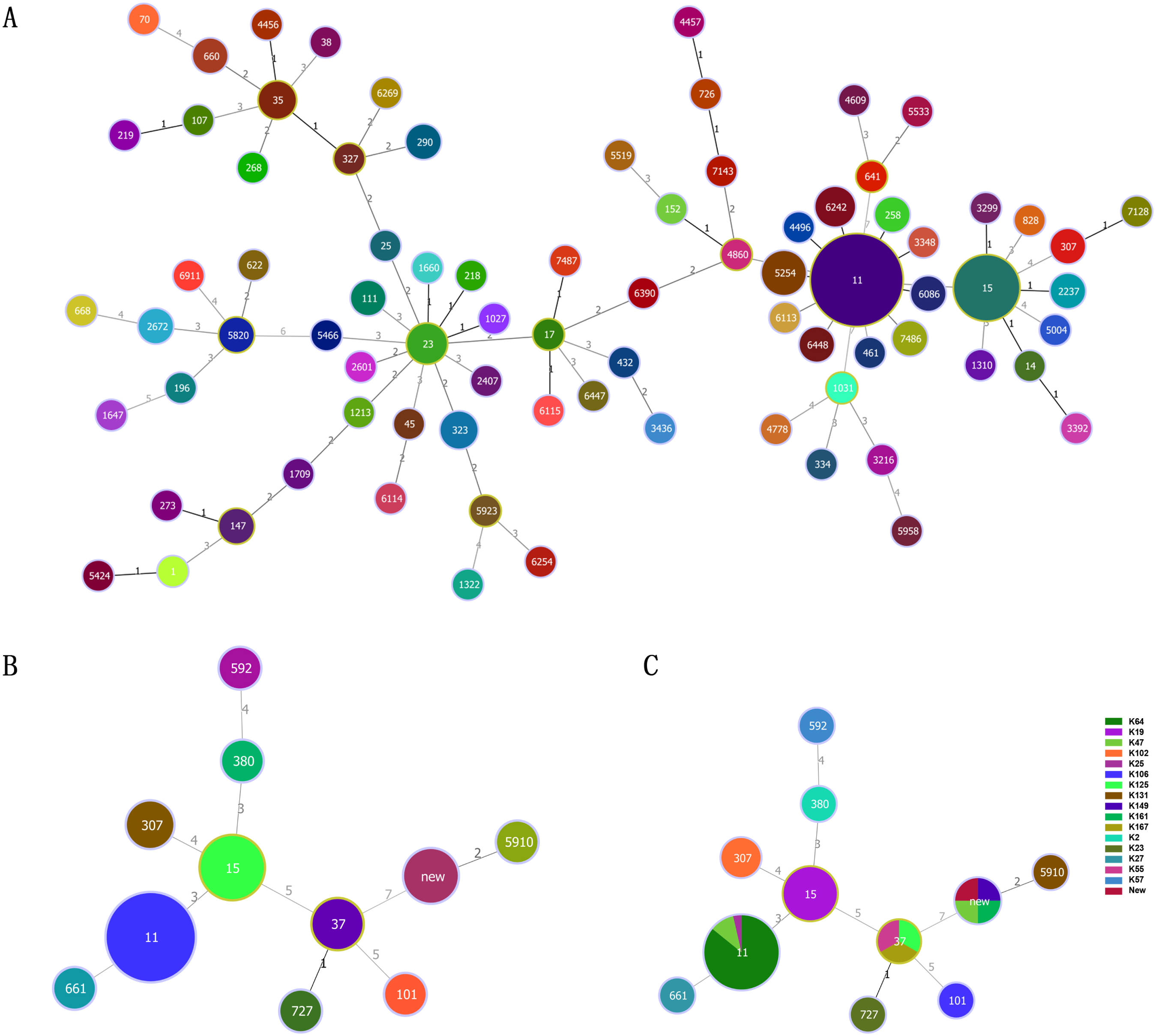
ST-type minimum-spanning tree. A.Genetic relationship of carbapenem-resistant *klebsiella pneumoniae* isolates of China. B. Genetic relationship of carbapenem-resistant *Klebsiella pneumonia* isolates of this study;C. Genetic relationship between MLST and K-type of carbapenem-resistant *Klebsiella pneumonia* isolates of this study. The size of the dots is proportional to the number of strains. Numbers on the connecting line represent genetic distance. ST, sequence typing

**Figure 4.**
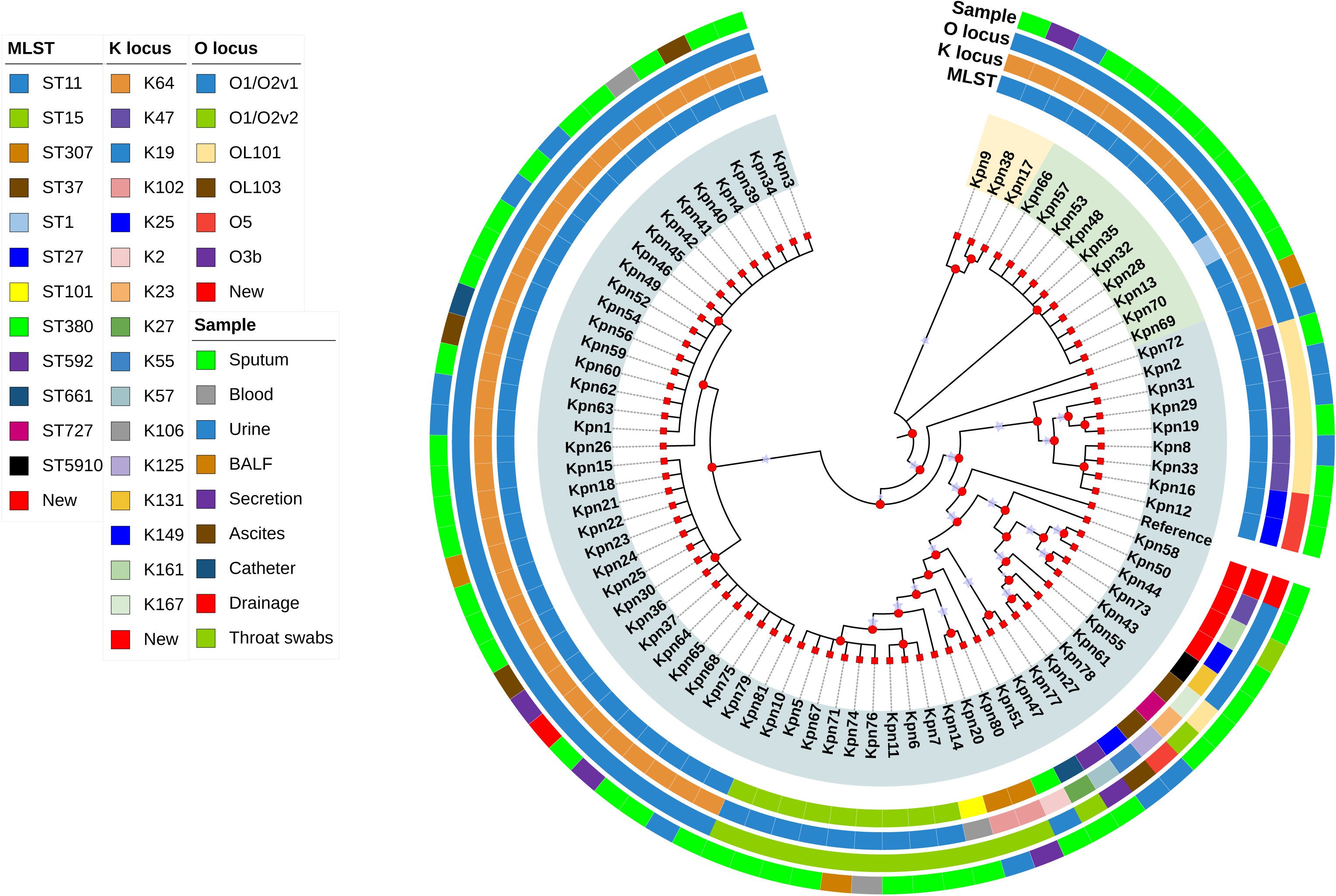
Evolutionary relationship diagram of 81 CRKP strains CRKP: carbapenem-resistant *Klebsiella pneumonia*

**Figure 5.**
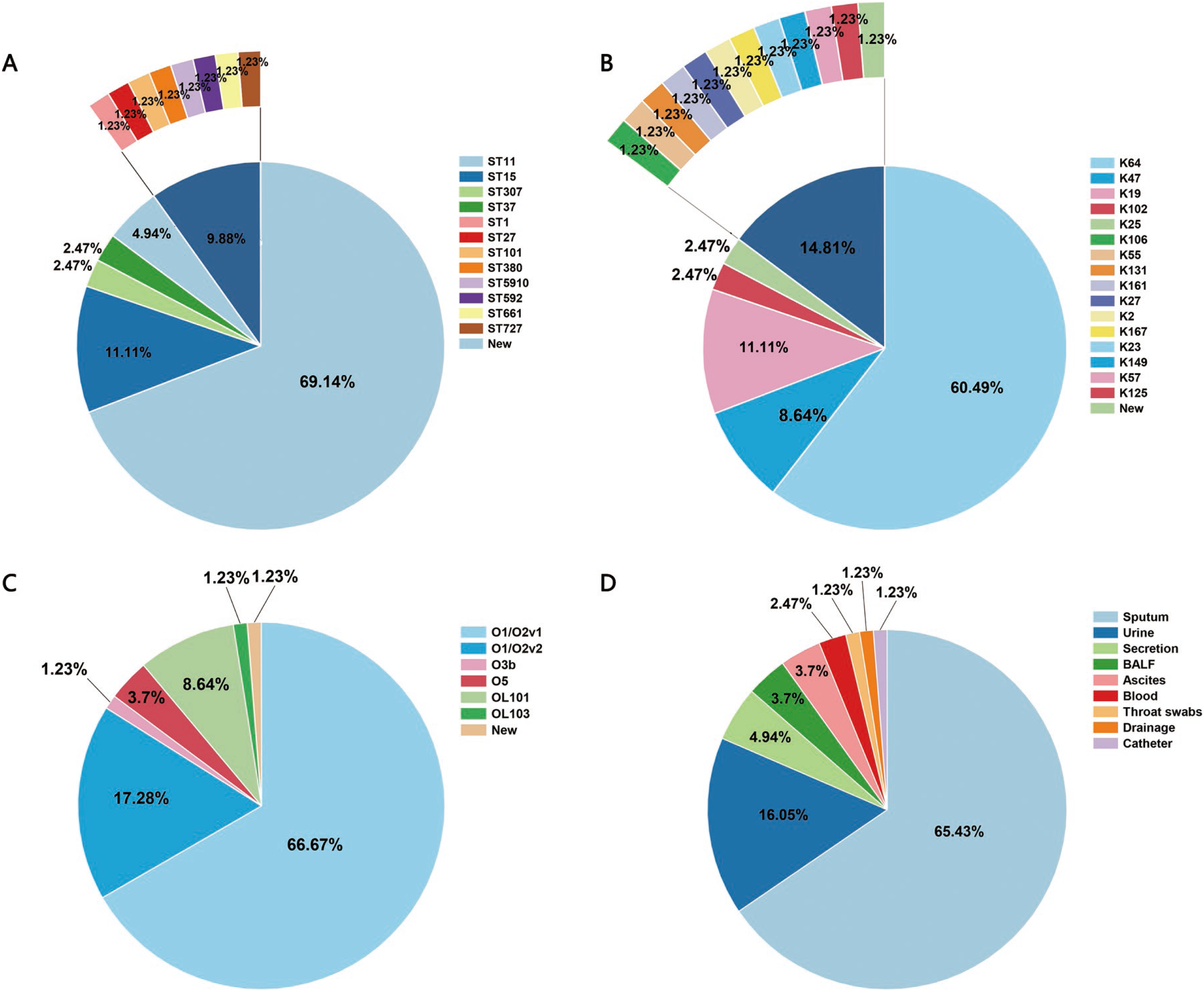
Overall distribution of 81 CRKP strains A. ST distribution; B. K-type distribution; C. O-type distribution; D. Sample distribution CRKP: carbapenem-resistant *Klebsiella pneumonia;* ST, sequence typing

### 2. Multivariate logistic regression analysis of independent risk factors for CR-hvKP infection

For the convenience of analysis, we categorized 81 CRKP strains into the following two groups: the high virulence (rmpA, rmpA2, iucA, iroB, and peg-344) (16) and low virulence groups. We collected clinical data from 81 patients with CRKP, including patient Basic Data, Department, Underlying diseases, Infection type, Invasive procedures and devices, Antimicrobial exposure, and Outcomes. After analysis, significant differences (p<0.05) were found between the two groups of patients regarding Antimicrobial usage time, Pulmonary disease, Malignant tumors, and Carbapenem antimicrobial exposure and Outcomes. Logistic multiple regression analysis was conducted on the items with significant differences mentioned above, and it was found that Antibiotic usage time, Carbapenem antibiotic exposure, and Malignant tumors were independent risk factors for CR-hvKP infection (Tables 1 and 2).

**Table 1.**
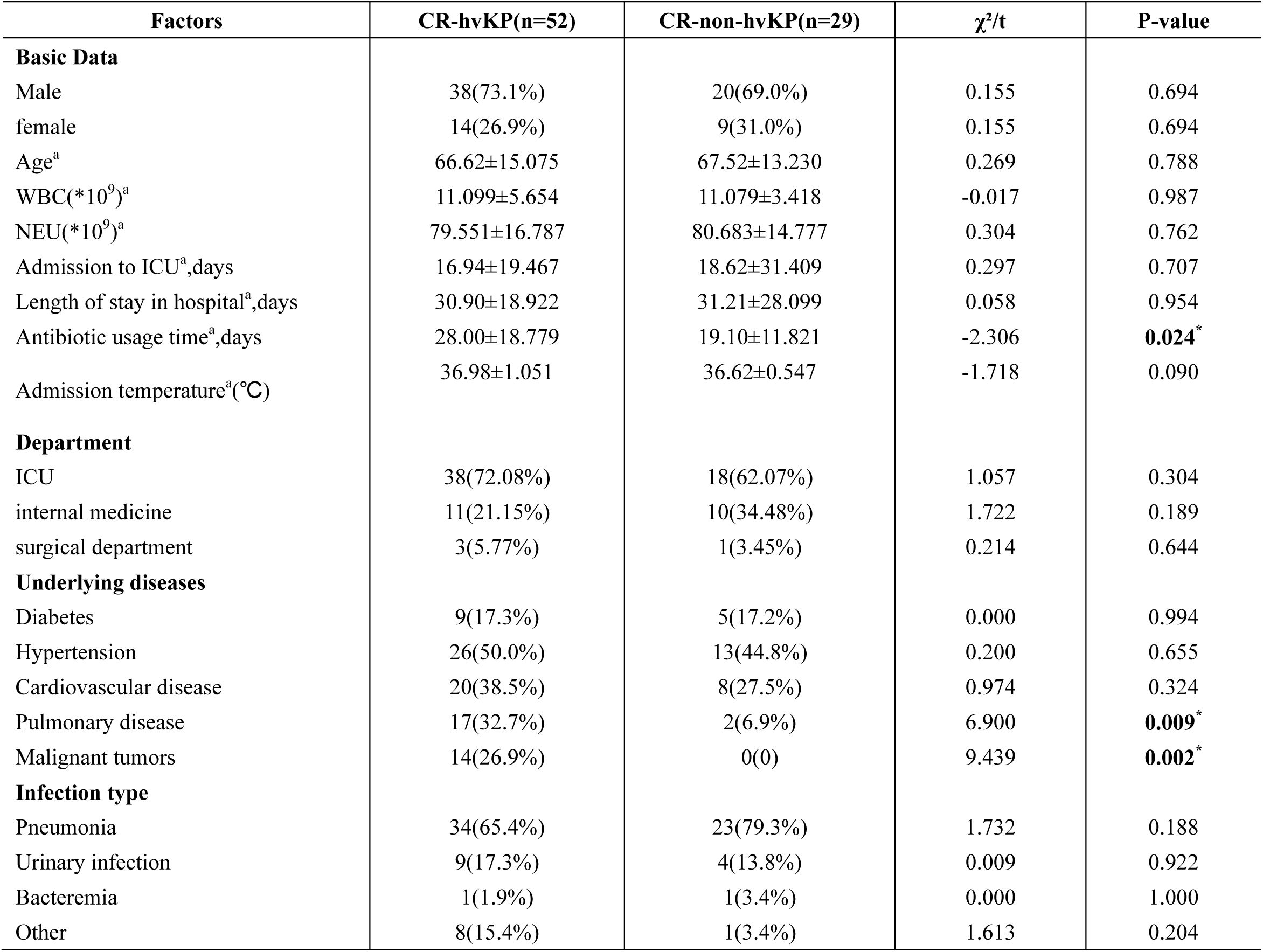
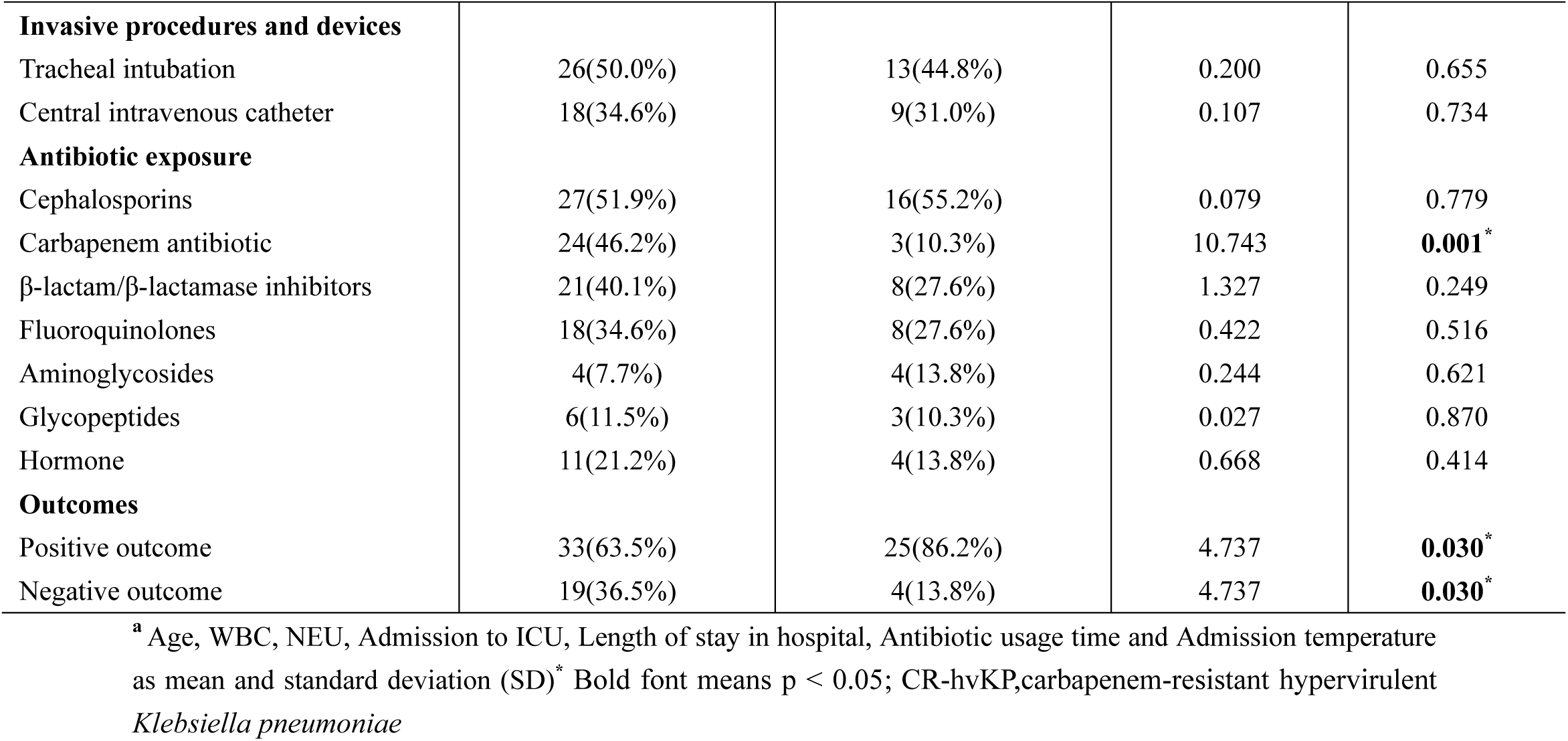
Clinical characteristics of CR-hvKP and CR-non-hvKP.

**Table 2.** Multivariate logistic regression analysis of independent risk factors for CR-hvKP infection.

| Variable | Univariate OR (95% CI) | p-value | Multivariate OR (95% CI) | p-value |
| --- | --- | --- | --- | --- |
| Antibiotic usage time | 1.042[1.008-1.083] | 0.0261 | 1.076[1.026-1.138] | **0.0051\*** |
| Carbapenem antibiotic | 0.3571[0.1341-0.9282] | 0.0358 | 0.117 [0.02266-0.4602] | **0.0045\*** |
| Pneumonia disease | 0.2975 [0.0995-0.8518] | 0.0254 | 1.594[0.8704-3.336] | 0.1593 |
| Malignant tumors | 13.6[2.546-252.4] | 0.0138 | 65.1[7.078-1798] | **0.0019\*** |
| Outcomes | 3.167[1.339-8.694] | 0.0138 | 0.2459[0.05379-0.9333] | 0.0504 |

\* Bold font means p < 0.05; OR, Odds Ratio; CR-hvKP, carbapenem-resistant hypervirulent *Klebsiella pneumoniae*

### 3. Antimicrobial susceptibility tests

We conducted a new antibiotics combination β-lactam/β-lactase inhibitor, tigecycline and polymyxin susceptibility test on 81 CRKP strains, including Aztreonam/Avibactam, Ceftaroline/Avibactam, Ceftazidim/Avibactam, Imipenem/Avibactam, Meropenem/Avibactam, and Etapenem/Avibatanm, and classified them into high (4 μg/mL) and low (8 μg/mL) concentration inhibitor groups. Except for Ceftaroline, significant differences (p<0.05) were found in the drug sensitivity results of the other five antibiotics after adding low-concentration avibactam inhibitors. The MIC values of Ceftazidim/Avibactam, Meropenem/Avibactam, and Etapenem/Avibatan increased as the inhibitor concentration rose (p<0.05). However, no significant change was observed in MIC values after adding high-concentration inhibitors, including Aztreonam/Avibactam and Imipenem/Avibactam (Figure 6). The resistance rate of polymyxin (79.01%) was lower than that of tigecycline (96.30%), and the resistance rate and MIC range values of the CR-hvKP group were generally higher than those of the CR-non-hvKP group (Tables 3 and 4).

**Figure 6.**
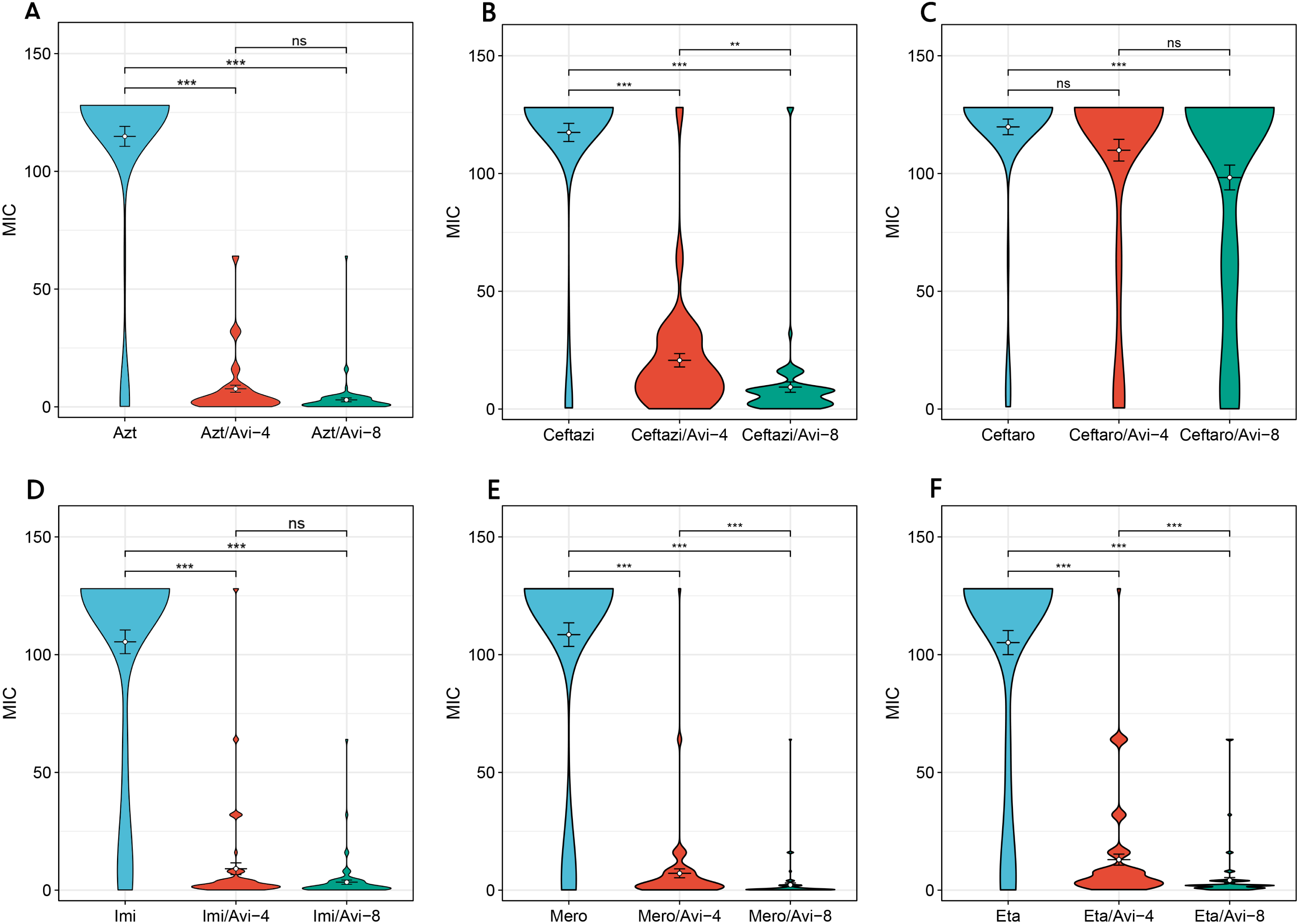
Comparison of MIC values of CRKP strains against novel antibiotics β-lactam/β-lactase inhibitors A.MIC values of Aztreonam, Aztreonam/Avibactam-4, and Aztreonam/Avibactam-8; B. MIC values of Ceftazidim, Ceftazidim/Avibactam-4, and Ceftazidim/Avibactam-8; C. MIC values of Ceftaroline, Ceftaroline/Avibactam-4, and Ceftaroline/Avibactam-8; D. MIC values of Imipenem, Imipenem/Avibactam-4, and Imipenem/Avibactam-8; E. MIC values of Mero penem, Meropenem/Avibactam-4, and Meropenem/Avibactam-8; F. MIC values of Etapene m, Etapenem/Avibatan-4, and Etapenem/Avibatan-8. “*” represents the size of P value, ns represents p≥0.05, *represents p<0.05, **represents p<0.01, and ***represents P<0.001. CRKP: carbapenem-resistant *Klebsiella pneumonia*; MIC: minimum inhibitory concentration; ST, sequence typing;Azt, Aztreonam;Ceftazi, Ceftazidime;Ceftaro, Ceftaroline;Imi, Imipenem;Mero, Meropenem;Eta, Ertapenem;Avi, Avibactam

**Table 3.** Drug susceptibility characteristics of novel enzyme inhibitor antibiotics.

| **Antibiotic** | **R** | **S** | **I** | **MIC** | **MIC50** | **MIC90** |
| --- | --- | --- | --- | --- | --- | --- |
| **Aztreonam** | 91.36% | 8.64% | 0.00% | >128 | >128 | >128 |
| **Aztreonam /Avibactam-4** | 13.58% | 72.84% | 13.58% | 4 | 1 | 2 |
| **Aztreonam /Avibactam-8** | 3.70% | 95.06% | 1.23% | 1 | 0.5 | 0.5 |
| **Ceftazidime** | 92.59% | 7.41% | 0.00% | >128 | >128 | >128 |
| **Ceftazidime/Avibactam-4** | 58.02% | 41.98% | 0.00% | 16 | 4 | 8 |
| **Ceftazidime/Avibactam-8** | 14.81% | 85.19% | 0.00% | 8 | 2 | 4 |
| **Ceftaroline** | 98.77% | 0.00% | 1.23% | >128 | >128 | >128 |
| **Ceftaroline/Avibactam-4** | 96.30% | 3.70% | 0.00% | >128 | >128 | >128 |
| **Ceftaroline/Avibactam-8** | 92.59% | 4.94% | 2.47% | >128 | 64 | 64 |
| **Imipenem** | 91.36% | 7.41% | 1.23% | >128 | >128 | >128 |
| **Imipenem/Avibactam-4** | 38.27% | 30.86% | 30.86% | 2 | 0.5 | 1 |
| **Imipenem/Avibactam-8** | 29.63% | 53.09% | 17.28% | 1 | 0.5 | 0.5 |
| **Meropenem** | 88.89% | 11.11% | 0.00% | >128 | >128 | >128 |
| **Meropenem/Avibactam-4** | 45.68% | 35.80% | 18.52% | 2 | 1 | 1 |
| **Meropenem/Avibactam-8** | 11.11% | 75.31% | 13.58% | 0.5 | <0.125 | 0.25 |
| **Ertapenem** | 91.36% | 7.41% | 1.23% | >128 | >128 | >128 |
| **Etapenem/Avibatan-4** | 85.19% | 7.41% | 7.41% | 4 | 2 | 2 |
| **Etapenem/Avibatan-8** | 56.79% | 19.75% | 23.46% | 2 | 0.5 | 1 |
| **Tigecycline** | 96.30% | 3.70% | 0.00% | 4 | 1 | 2 |
| **Polymyxin** | 79.01% | 20.99% | 0.00% | 8 | 2 | 4 |

R, drug resistance rate ; S, sensitivity rate; I , Intermediary rate

**Table 4.** Resistance rates of CR-hvKP and CR-non-hvKP.

| Antibiotic | CR-hvKP |  | CR-non-hvKP |  |
| --- | --- | --- | --- | --- |
|  | R% | MIC range | R% | MIC range |
| **Aztreonam** | 98.08% | 0.25->128 | 79.31% | 0.25->128 |
| **Aztreonam /Avibactam-4** | 17.31% | 0.125-64 | 6.90% | <0.125-64 |
| **Aztreonam /Avibactam-8** | 1.92% | <0.125-64 | 6.90% | <0.125-16 |
| **Ceftazidime** | 98.08% | 0.5->128 | 82.76% | 0.5->128 |
| **Ceftazidime/Avibactam-4** | 71.15% | 1-64 | 34.48% | <0.125->128 |
| **Ceftazidime/Avibactam-8** | 13.46% | 0.5-16 | 17.24% | <0.125->128 |
| **Ceftaroline** | 100.00% | 16->128 | 96.55% | 1->128 |
| **Ceftaroline/Avibactam-4** | 100.00% | 4->128 | 89.66% | 0.5->128 |
| **Ceftaroline/Avibactam-8** | 100.00% | 4->128 | 79.31% | <0.125->128 |
| **Imipenem** | 96.15% | <0.125->128 | 82.76% | 0.5-64 |
| **Imipenem/Avibactam-4** | 46.15% | 0.25-64 | 24.14% | <0.125->128 |
| **Imipenem/Avibactam-8** | 32.69% | <0.125-64 | 24.14% | <0.125->128 |
| **Meropenem** | 100.00% | 16->128 | 68.97% | <0.125->128 |
| **Meropenem/Avibactam-4** | 53.85% | 0.125-16 | 31.03% | <0.125->128 |
| **Meropenem/Avibactam-8** | 9.62% | <0.125-16 | 13.79% | <0.125->128 |
| **Ertapenem** | 98.08% | 0.25->128 | 79.31% | <0.125-64 |
| **Etapenem/Avibatan-4** | 96.15% | 0.25-64 | 65.52% | <0.125-128 |
| **Etapenem/Avibatan-8** | 69.23% | <0.125-64 | 34.48% | <0.125-64 |
| **Tigecycline** | 96.15% | 0.5-64 | 96.55% | 0.25-128 |
| **Polymyxin** | 86.54% | <0.125-128 | 65.52% | <0.125-64 |
| **Cefepime** | 88.46% | >16->32 | 100.00% | <1->32 |
| **Cefotaxime** | 80.77% | >2->32 | 66.00% | <0.5->32 |
| **Amoxicillin/clavulanate** | 88.46% | >16/8 | 100.00% | <4/2->16/8 |
| **Ampicillin/Sulbactam** | 96.15% | >16/8 | 100.00% | 8/4->16/8 |
| **Levofloxacin** | 95.83% | >4->8 | 100.00% | <1->8 |
| **Ciprofloxacin** | 92.31% | >2->16 | 100.00% | <1->16 |
| **Amikacin** | 23.08% | <8->32 | 92.00% | <8->32 |
| **Gentamicin** | 73.08% | <2->8 | 90.00% | <1->8 |

\* Bold font means p < 0.05; R, drug resistance rate; CR-hvKP, carbapenem-resistant hypervirulent *Klebsiella*
*pneumoniae*

### 4. Antimicrobial resistance genes, virulence genes, and plasmids of CR-hvKP

All 52 CR-hvKP strains carried the carbapenemase gene bla_KPC-2_. Carbapenem resistance genes included bla_KPC-2_ and bla_OXA-1_. Chloramphenicol resistance genes were found only in the ST11-K47-OL101 strain. The bla_CTX-M-15_ gene was detected only in ST307-K102-O1/O2v2, while SHV-182 was found in both ST307-K102-O1/O2v2 and ST1-K64-O1/O2v1. All strains carry bla_KPC-2_, ompk35, and ompk36 genes (Figure 7 and Figure 8A). Carbapenem resistance genes bla_OXA-1_ and bla_CTX-M-15_ were strongly correlated with IncFII_1_pKP91, while tetracycline resistance genes were strongly associated with IncFII(pCRY) (Figure 9B). The resistance genes of macrolides antibiotics were positively correlated with CRP (Figure 9D).

**Figure 7.**
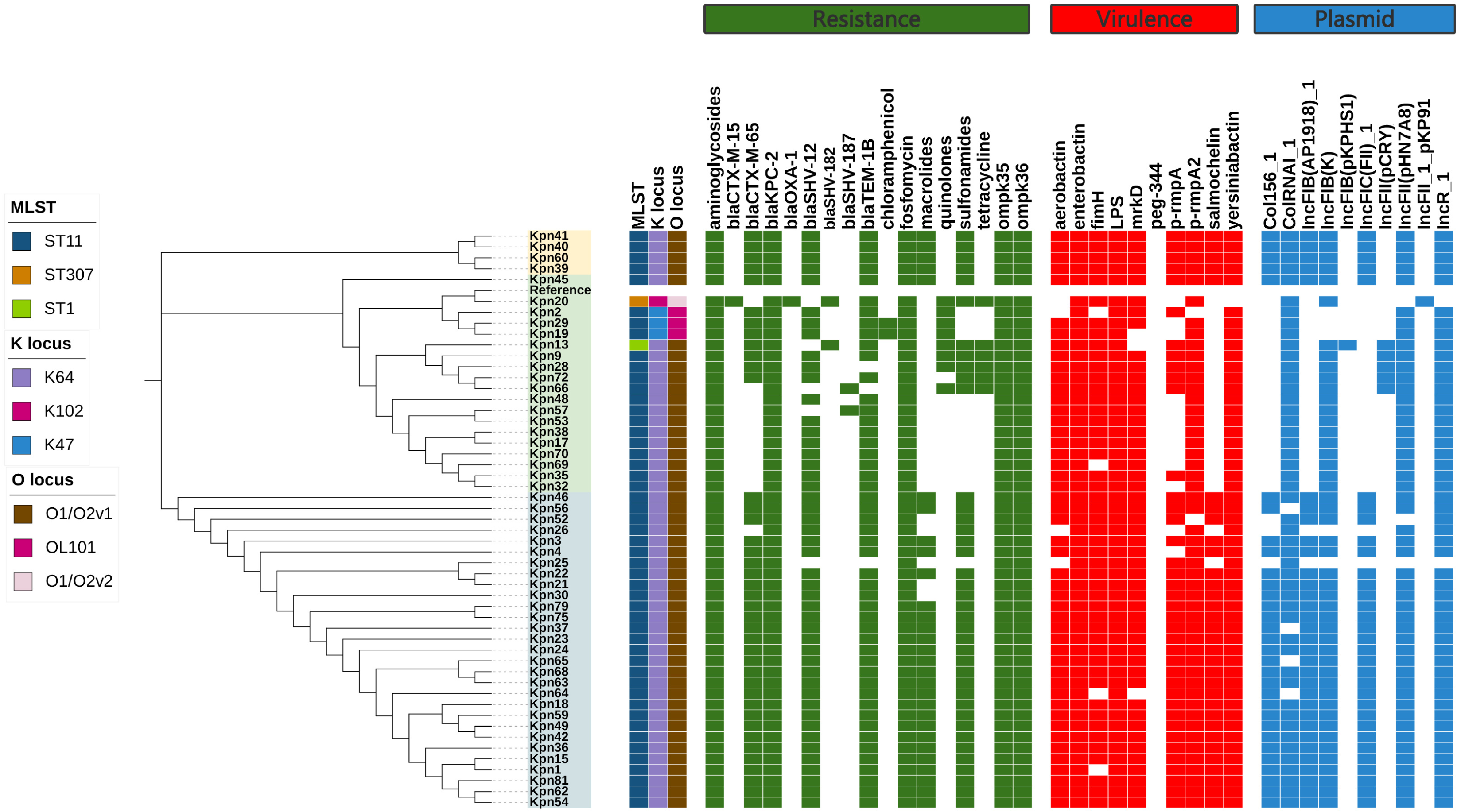
Evolutionary relationship diagram of 52 CR-hvKP strains. CR-hvKP: carbapenem-resistant hypervirulent *Klebsiella pneumoniae*

**Figure 8.**
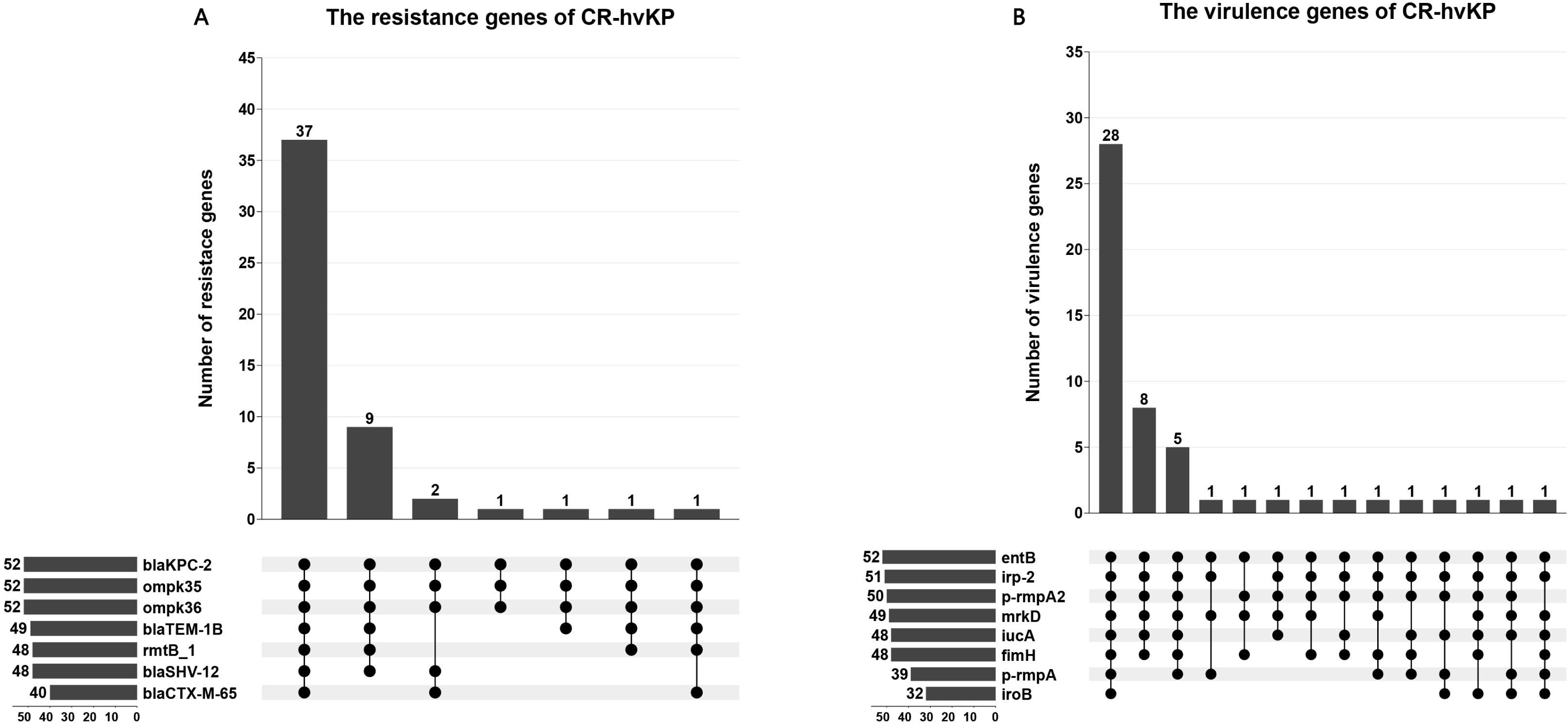
The characteristic distribution of resistance and virulence genes. A. Distribution of resistance genes; B. Distribution of virulence genes.

**Figure 9.**
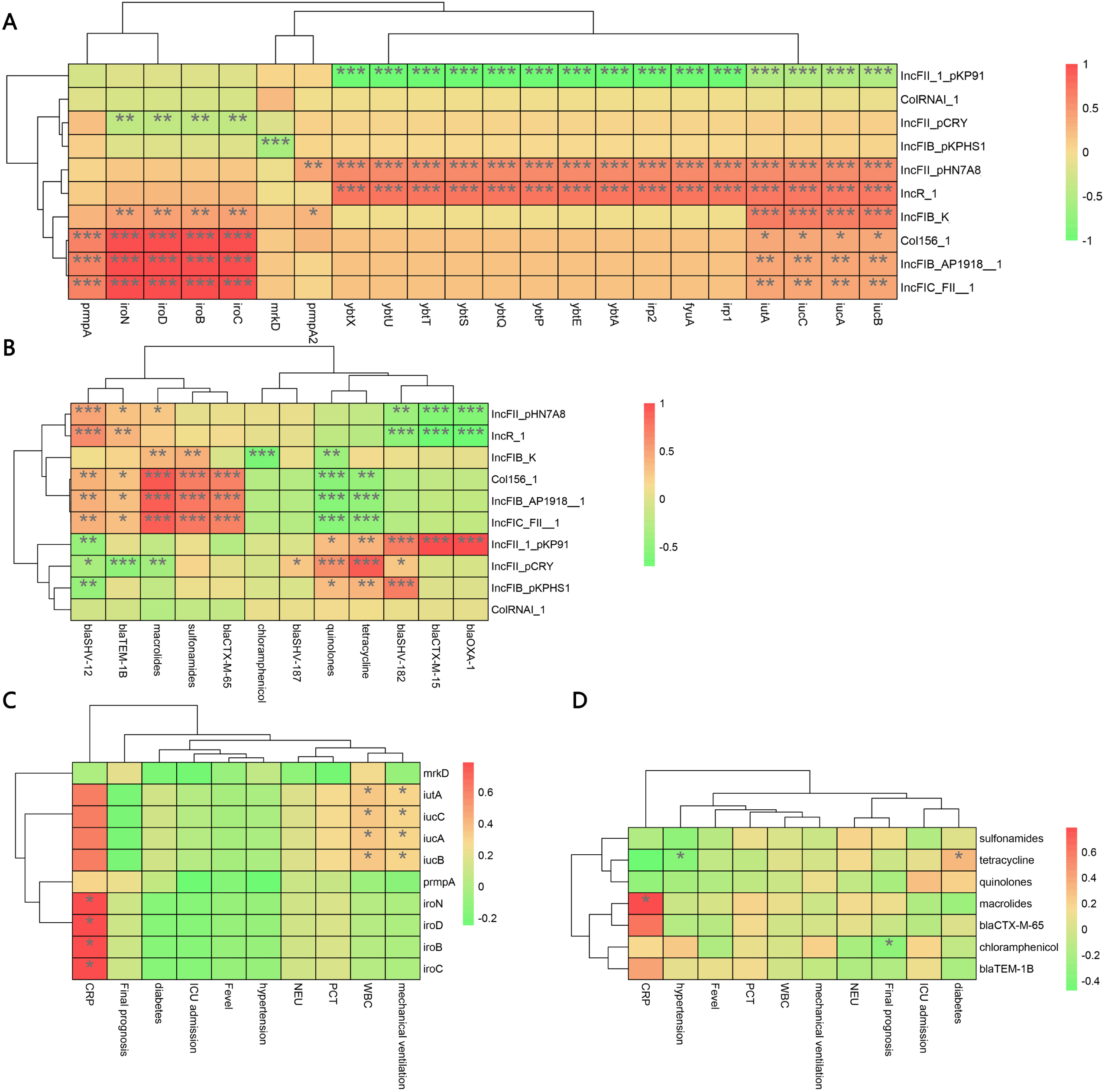
Correlations among virulence genes, resistance genes, plasmid replicons, and clinical data of CR-hvKP strain. A. Spearman correlation between virulence genes and plasmid replicon. B. Spearman correlation between resistance genes and plasmid replicon. C. Spearman correlation between virulence genes and clinical data. D. Spearman correlation between resistance genes and clinical data. “*” represents the size of the P value, ns represents p≥0.05, *represents p<0.05, **represents p<0.01, and ***represents p<0.001. CR-hvKP: carbapenem-resistant hypervirulent Klebsiella pneumoniae

Virulence gene analysis revealed that 28 CR-hvKP strains carried all tested virulence genes. The lowest detection rates were observed for iroB (61.54%) (Figure 8B). Siderophore virulence genes showed high carrying rates in all strains. Salmonella was detected in 61.54% of the strains, which was lower than that for *aerobactin* (92.31%), *yersiniabactin* (98.08%), and *enterobactericin* (100%). The *yersiniabactin* gene was not detected in ST307-K102-O1/O2v2. No strains tested positive for peg-344 (metabolite transporter) virulence genes. Among the 52 CR-hvKP strains, 10 plasmid replicon types were identified (Figure 7). *Aerobactin* and *yersiniabactin* virulence genes were strongly correlated with the IncR_1 (Figure 9A). The virulence genes of Salmochelin were positively correlated with CRP (Figure 9C).

The experiment on the *Galleria mellonella* showed that the survival rate of CR-non-hvKP was higher than that of CR-hvKP. The survival rate of CR-hvKP-47 was higher than that of CR-hvKP-64, indicating that the virulence of CR-hvKP-64 was higher than that of CR-hvKP-47 (Figure 10).

**Figure 10.**
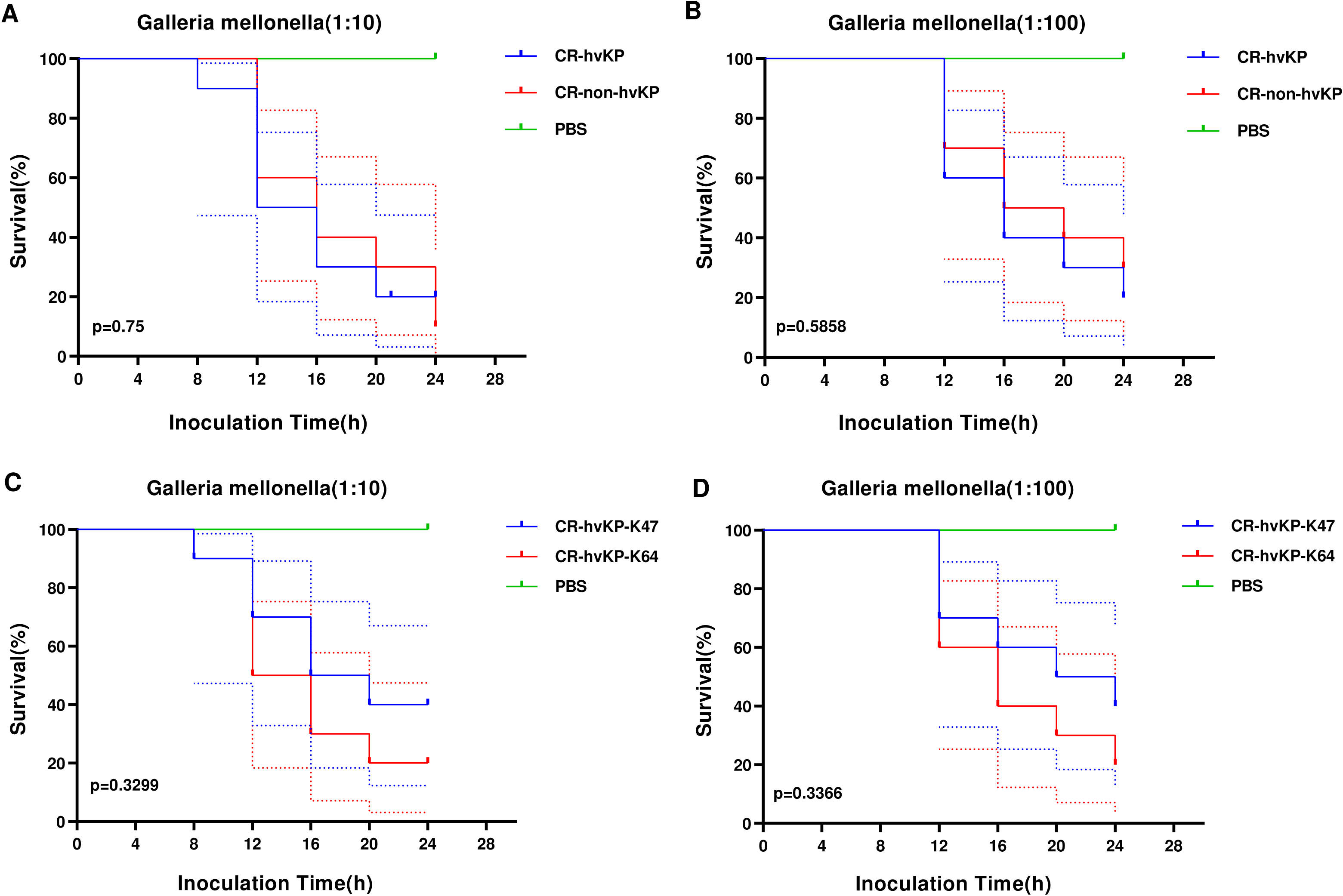
The *Galleria mellonella* infection model. A. Survival rates of *Galleria mellonella* infected (1:10) with CR-hvKP and CR-non-hvKP; B. Survival rates of *Galleria mellonella* infected (1:100) with CR-hvKP and CR-non-hvK P; C. Survival rates of *Galleria mellonella* infected (1:10) with CR-hvKP-K47 and CR-h vKP-K64; D. Survival rates of *Galleria mellonella* infected (1:100) with CR-hvKP-K47 and CR-hvKP-K64

### 5. Antibiotic resistance plasmids and mobile genetic elements

Blast analysis showed that these drug-resistant plasmids were highly similar to the reference plasmid pC76 KPC (NZ_CP080299.1) with 99.37% identity and 93.25% coverage. The plasmid does not contain the fusion plasmid of virulence, non-drug resistance and virulence genes. pKpn30, pKpn45, and pKpn46 were similar in structure and belonged to the plasmid replicons of IncFIB(AP001918)_1, IncFIC(FII)_1, and IncFII(pHN7A8)_1. The structures of pKpn32, pKpn38, and pKpn70 were similar, belonging to repB_KLEB_VIR, IncHI1B(pNDM-MAR)_1, and IncFII(pHN7A8)_1 plasmid replicons. After analysis, the bla_KPC-2_ genes of the six CR-hvKP strains were all located on plasmids and carried bla_TEM-1B_, bla_SHV-12,_ and rmtB_1 resistance genes. Various transposons and insertion sequences around the resistance genes predicted the possibility of horizontal transfer of the resistance genes. Further study of the genetic environment of bal_KPC-2_ genes revealed that the upstream and downstream of the bal_KPC-2_ gene integrated TnpR_Tn3, ISKpn27, and ISKpn6. ISKpn27 contained IRL and IRR, while ISKpn6 only had IRR (Figures 11 and 12).

**Figure 11.**
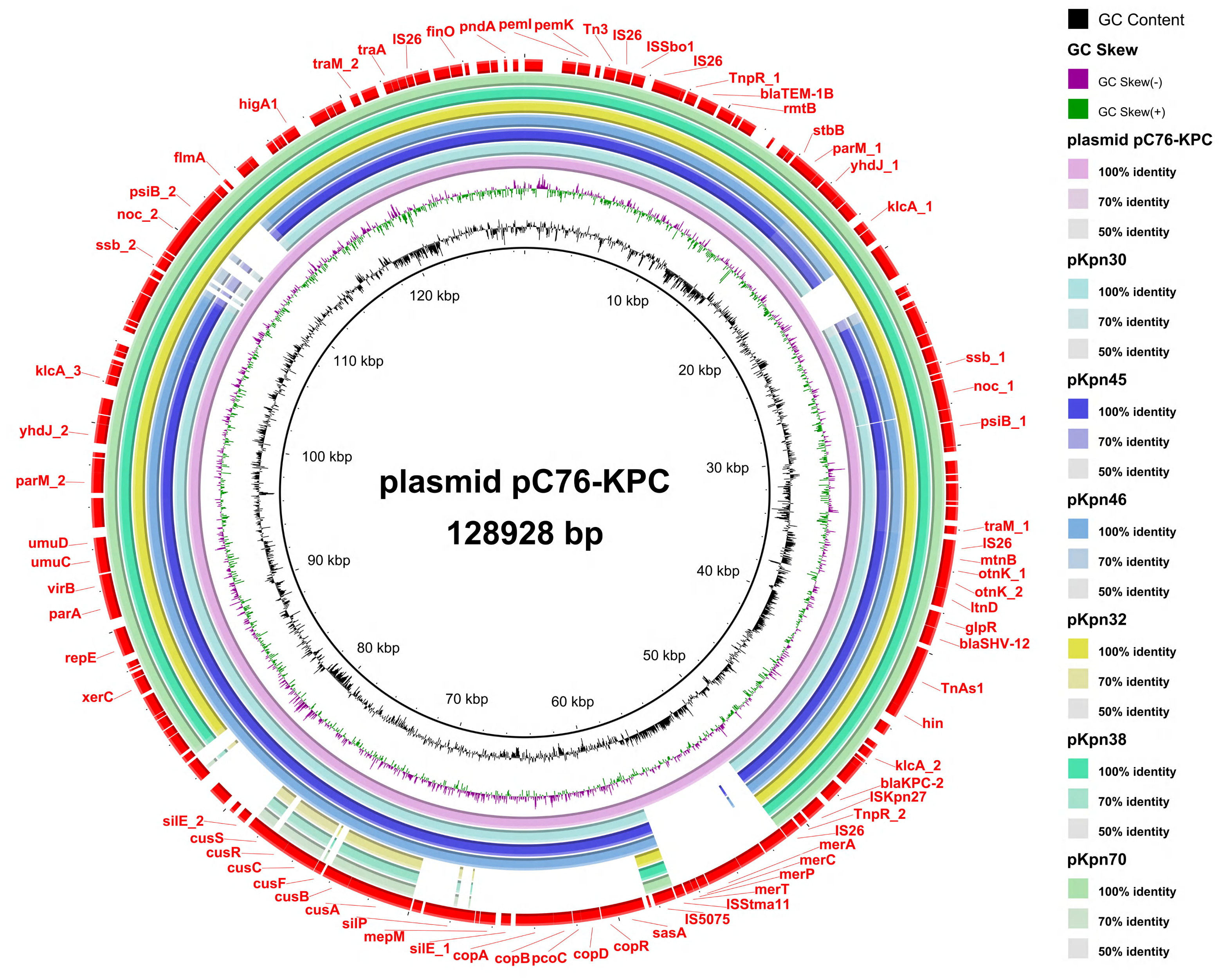
Comparative genomic analysis of antibiotic-resistant plasmids in CR-hvKP strains. From the inner circle to the outer circle, use this as follows: circle 1, GC Content; circle 2, GC Skew; circle 3, reference plasmid pC76 KPC(NZ_CP080299.1); circle 4, pKpn30; circle 5, pKpn45; circle 6, pKpn46; circle 7, pKpn32; circle 8, pKpn38; circle 9, pKpn 70; and circle 10, Annotated mobile genetic elements and carbapenem resistance genes. CR-hvKP: carbapenem-resistant hypervirulent *Klebsiella pneumoniae*

**Figure 12.**
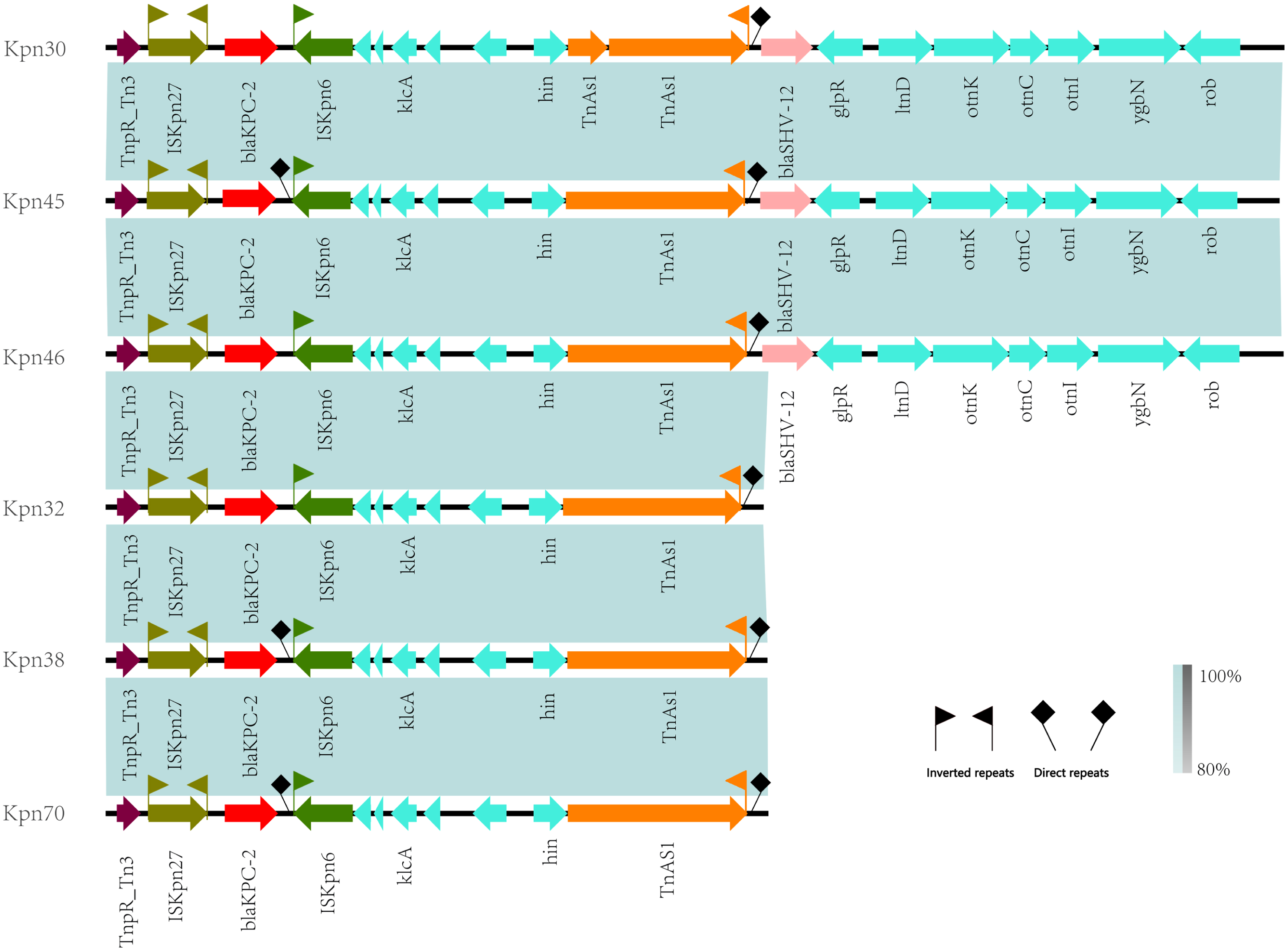
Linear alignment of gene environment in bla_KPC-2_.

## Discussion

Recently, the prevalence of CR-hvKP has been on the rise globally (17). According to the China Antimicrobial Surveillance Network, the resistance rate of *KP* to carbapenems has increased approximately ninefold, from 2.95% in 2005 to 25.4% in 2023, making it the second most commonly detected bacterium after *Escherichia coli* (https://www.chinet.gov/Data/GermYear). Similarly, data from the European Center for Disease Prevention and Control show an increase in carbapenem resistance in *KP* from 7.1% in 2017 to 11.7% in 2021 (https://www.ecdc.europa.eu/en). The high pathogenicity and resistance of CR-hvKP frequently lead to poor clinical outcomes, posing significant challenges for infection control in healthcare settings owing to its rapid spread and outbreaks. Studies have demonstrated that underlying chronic comorbidities, diabetes, age <65 years, and mechanical ventilation are risk factors for CR-hvKP infections (18, 19).

In our hospital, the predominant MLST is ST11, consistent with the findings of previous studies by Cao et al. (5). In contrast, ST258 is the major circulating type in Europe and the United States (20, 21). Recent research from Turkey has identified ST101 as the most common type among CRKP infection strains (22), while a study on mother-child cohorts in Madagascar found that ST23 and ST25 are associated with high virulence (23). In Argentina, high-mucoviscosity CRKP strains were primarily isolated from respiratory and urinary samples (34.2%), with 40% of strains belonging to ST25 and all strains carrying the bla_KPC-2_ resistance gene (24). In Greece, bloodstream infections are the primary type of CRKP infections, and ST39 carrying bla_KPC-2_ is considered a high-risk clone (4). Regions such as Egypt, India, Pakistan, and Serbia are major hotspots for the spread of the Class B carbapenemase NDM (25). The Class D carbapenemase OXA-48 has lower hydrolytic activity but frequently combines with other resistance mechanisms to increase its effectiveness. The spread of bla_OXA-232_ ST15 *KP* clones has occurred globally, particularly in China, although they likely originated in the United States (26). However, the emergence of CRKP strains that produce more than one type of carbapenemase is a concern (27). In our study, one high-risk clone, ST307, contained no *yersiniabactin*, *aerobactin*, or *salmochelin*. Some studies have demonstrated that ST307 *KP* can increase the MIC value of ceftazidime/avibactam, leading to resistance (28). In some regions, ST307 has replaced the high-risk clones ST512 and ST258 (29). We should be vigilant about the emergence of ST307 *KP* and strengthen monitoring of this clone type promptly. It has been discovered that *KP* contains at least 79 K-capsule types. In China, the most common K types of CRKP are K64 (50.4%) and K47 (25.9%). The detection rate of K64 has been increasing, mainly in eastern and central China, whereas that of K47 has significantly declined and is primarily found in northern and northeastern China (3). Hu et al. indicated that capsule-deficient strains have defects in transmission and resistance to phagocytosis; however, their antibiotic resistance increases correspondingly (3). This was confirmed in previous studies, which reported that capsule-deficient strains cause untreatable persistent urinary tract infections, whereas hypercapsular strains showed enhanced resistance to phagocytosis, increased dissemination, and higher mortality rates (21). The rising prevalence of CRKP and changes in its serotypes suggest an urgent need to develop vaccines against CRKP infections. Our study found that the virulence of CR-hvKP-K64 was higher than that of K47-CR-hvKP, which is consistent with the findings of Jia et al. (30).

Avibactam is a β-lactamase inhibitor that can inhibit serine protease activity but is ineffective against metalloenzyme activity. Owing to its excellent *in vitro* efficacy and safety against serine carbapenemases, ceftazidime/avibactam has become the first-line treatment for CRKP (31). However, the emergence of bla_KPC_ mutations, including bla_KPC-135_ and bla_KPC-112_, which are resistant to ceftazidime/avibactam, has been reported due to the improper use of antibiotics (6, 32). The spread of bla_KPC_ can be mediated through various molecular mechanisms, including small genetic elements, horizontal transfer of plasmids, and clonal transmission. In our study, Ceftaroline/Avibactam resistance is mainly mediated by the plasmid pC76 KPC (NZ_CP080299.1) carrying bla_KPC-2_. This plasmid is nonclassical, and it was first included in the RefSeq microbial genomes database in the United States (33). The pKpQIL plasmid was the first identified from the ST258 strain carrying bla_KPC_, while the pQP048 plasmid, widely prevalent in China, carries bla_KPC_ and is most commonly associated with the ST11 strain (7). Mobile elements play a crucial role in the transfer of bla_KPC_ genes to different plasmids. The most common mobile element containing bla_KPC_ is the Tn4401 transposon based on Tn3, predominantly found in the United States, while NTEXPC-I and NTEXPC-II are primarily distributed in China and Brazil (34). Further research on the genetic environment of the bla_KPC-2_ gene revealed that the upstream and downstream regions of bla_KPC-2_ genes contain insertion sequences of ISKpn27 and ISKpn6, respectively. There are incomplete reverse repeat sequences on both sides of ISKpn27, which makes the bla_KPC-2_ gene easy to spread. Current research has found that various bla_KPC-2_ subtypes are associated with distinct mobile genetic elements and located in different plasmids. Notably, a study found that a non-ribosomal tobramycin-cyclohexane conjugate could enhance the efficacy of β-lactam/β-lactamase inhibitors against carbapenem-resistant clinical isolates (35), providing a new approach to maintaining the therapeutic utility of β-lactamase antibiotics. Our study showed that the rate of resistance to polymyxins was lower than that to tigecycline. Yu et al. demonstrated that hypervirulent ST11-K64 could develop resistance through rapid and diverse mechanisms during tigecycline and polymyxin treatment (36). Additionally, the emergence of tmexCD-toprJ has significantly reduced the effectiveness of tigecycline against CRKP (37). Recently, increasing reports of polymyxin-resistant *KP* have highlighted a major challenge to public health owing to the emergence of colistin-resistant and hvKP (38).

The limitations of this study include its single-center data, which only represents the epidemiological characteristics of CRKP in this specific region. In future research, we plan to use third-generation genome sequencing for a comprehensive analysis of whole-genome sequences and continue expanding the data collection to complete a multi-center data analysis.

## Conclusion

In this study, we conducted a systematic molecular epidemiological analysis of CRK P. The bla_KPC-2_ gene was identified as the primary mechanism of carbapenem resistance in our hospital, with ST11-K64 emerging as the dominant clone. The drug-resistant plasmid pC76-KPC(NZ_CP080299.1) containing the bla_KPC-2_ gene mediates Ceftazidim/Avibactam resistance. This study contributes to the optimization of antimicrobial management; moreover, genomic research can aid in tracking hospital infection outbreaks and improving infection control measures.

## Data availability statement

Data from this study can be available upon request from the author.

## Acknowledgments

We thank all colleagues and classmates in the laboratory for their hard work in the experiment. We also thank colleagues in the Microbiology Department of the First Affiliated Hospital of Hebei North University for their technical support. The author thanks Editage (www.editage.cn) for English language editing.

## Ethics declarations

### Ethics approval and consent to participate

The study was approved by the Ethics Committee of the First Affiliated Hospital of Hebei North University (ethical approval No. K2019147), which waived the requirement of written informed consent from patients. All strains are part of the routine laboratory procedures of the hospital and do not involve any human genetic resources. This study was conducted in accordance with the principles outlined in the Declaration of Helsinki.

### Consent for publication

Not applicable.

### Authors’ contributions

WZ was responsible for conceptualization. BW sequenced the bacterial strain. NW, MZ, LD, NJ, XP, JC, and JH conducted the experiments. HL, LS, TW, and LC data curation. CL and BL, formal analysis. NW, writing – original draft, and visualization. All authors agree to the final version of the article.

## References

1. Zhang Y, Wang Q, Yin Y, Chen H, Jin L, Gu B, et al. Epidemiology of Carbapenem-Resistant Enterobacteriaceae Infections: Report from the China CRE Network. Antimicrob Agents Chemother. 2018;62(2).

2. Yang X, Sun Q, Li J, Jiang Y, Li Y, Lin J, et al. Molecular epidemiology of carbapenem-resistant hypervirulent Klebsiella pneumoniae in China. Emerg Microbes Infect. 2022;11(1):841–9.

3. Hu F, Pan Y, Li H, Han R, Liu X, Ma R, et al. Carbapenem-resistant Klebsiella pneumoniae capsular types, antibiotic resistance and virulence factors in China: a longitudinal, multi-centre study. Nat Microbiol. 2024;9(3):814–29.

4. Tryfinopoulou K, Linkevicius M, Pappa O, Alm E, Karadimas K, Svartström O, et al. Emergence and persistent spread of carbapenemase-producing Klebsiella pneumoniae high-risk clones in Greek hospitals, 2013 to 2022. Euro Surveill. 2023;28(47).

5. Pu D, Zhao J, Lu B, Zhang Y, Wu Y, Li Z, et al. Within-host resistance evolution of a fatal ST11 hypervirulent carbapenem-resistant Klebsiella pneumoniae. Int J Antimicrob Agents. 2023;61(4):106747.

6. Shi Q, Shen S, Tang C, Ding L, Guo Y, Yang Y, et al. Molecular mechanisms responsible KPC-135-mediated resistance to ceftazidime-avibactam in ST11-K47 hypervirulent Klebsiella pneumoniae. Emerg Microbes Infect. 2024;13(1):2361007.

7. Yang X, Dong N, Chan EW, Zhang R, Chen S. Carbapenem Resistance-Encoding and Virulence-Encoding Conjugative Plasmids in Klebsiella pneumoniae. Trends Microbiol. 2021;29(1):65–83.

8. Gu D, Dong N, Zheng Z, Lin D, Huang M, Wang L, et al. A fatal outbreak of ST11 carbapenem-resistant hypervirulent Klebsiella pneumoniae in a Chinese hospital: a molecular epidemiological study. Lancet Infect Dis. 2018;18(1):37–46.

9. Pu D, Zhao J, Chang K, Zhuo X, Cao B. “Superbugs” with hypervirulence and carbapenem resistance in Klebsiella pneumoniae: the rise of such emerging nosocomial pathogens in China. Sci Bull (Beijing). 2023;68(21):2658–70.

10. Ribeiro-Gonçalves B, Francisco AP, Vaz C, Ramirez M, Carriço JA. PHYLOViZ Online: web-based tool for visualization, phylogenetic inference, analysis and sharing of minimum spanning trees. Nucleic Acids Res. 2016;44(W1):W246–51.

11. Letunic I, Bork P. Interactive Tree of Life (iTOL) v6: recent updates to the phylogenetic tree display and annotation tool. Nucleic Acids Res. 2024;52(W1):W78–w82.

12. Darling AC, Mau B, Blattner FR, Perna NT. Mauve: multiple alignment of conserved genomic sequence with rearrangements. Genome Res. 2004;14(7):1394–403.

13. Alikhan NF, Petty NK, Ben Zakour NL, Beatson SA. BLAST Ring Image Generator (BRIG): simple prokaryote genome comparisons. BMC Genomics. 2011;12:402.

14. Siguier P, Perochon J, Lestrade L, Mahillon J, Chandler M. ISfinder: the reference centre for bacterial insertion sequences. Nucleic Acids Res. 2006;34(Database issue):D32–6.

15. Sullivan MJ, Petty NK, Beatson SA. Easyfig: a genome comparison visualizer. Bioinformatics. 2011;27(7):1009–10.

16. Russo TA, Olson R, Fang CT, Stoesser N, Miller M, MacDonald U, et al. Identification of Biomarkers for Differentiation of Hypervirulent Klebsiella pneumoniae from Classical K. pneumoniae. J Clin Microbiol. 2018;56(9).

17. Han YL, Wen XH, Zhao W, Cao XS, Wen JX, Wang JR, et al. Epidemiological characteristics and molecular evolution mechanisms of carbapenem-resistant hypervirulent Klebsiella pneumoniae. Front Microbiol. 2022;13:1003783.

18. Liang S, Cao H, Ying F, Zhang C. Report of a Fatal Purulent Pericarditis Case Caused by ST11-K64 Carbapenem-Resistant Hypervirulent Klebsiella pneumoniae. Infect Drug Resist. 2022;15:4749–57.

19. Li L, Li S, Wei X, Lu Z, Qin X, Li M. Infection with Carbapenem-resistant Hypervirulent Klebsiella Pneumoniae: clinical, virulence and molecular epidemiological characteristics. Antimicrob Resist Infect Control. 2023;12(1):124.

20. David S, Reuter S, Harris SR, Glasner C, Feltwell T, Argimon S, et al. Epidemic of carbapenem-resistant Klebsiella pneumoniae in Europe is driven by nosocomial spread. Nat Microbiol. 2019;4(11):1919–29.

21. Ernst CM, Braxton JR, Rodriguez-Osorio CA, Zagieboylo AP, Li L, Pironti A, et al. Adaptive evolution of virulence and persistence in carbapenem-resistant Klebsiella pneumoniae. Nat Med. 2020;26(5):705–11.

22. Ibik YE, Ejder N, Sevim E, Rakici E, Tanriverdi ES, Copur Cicek A. Evaluating molecular epidemiology of carbapenem non-susceptible Klebsiella pneumoniae isolates with MLST, MALDI-TOF MS, PFGE. Ann Clin Microbiol Antimicrob. 2023;22(1):93.

23. Rakotondrasoa A, Passet V, Herindrainy P, Garin B, Kermorvant-Duchemin E, Delarocque-Astagneau E, et al. Characterization of Klebsiella pneumoniae isolates from a mother-child cohort in Madagascar. J Antimicrob Chemother. 2020;75(7):1736–46.

24. Vargas JM, Moreno Mochi MP, Nuñez JM, Cáceres M, Mochi S, Del Campo Moreno R, et al. Virulence factors and clinical patterns of multiple-clone hypermucoviscous KPC-2 producing K. pneumoniae. Heliyon. 2019;5(6):e01829.

25. Wu W, Feng Y, Tang G, Qiao F, McNally A, Zong Z. NDM Metallo-β-Lactamases and Their Bacterial Producers in Health Care Settings. Clin Microbiol Rev. 2019;32(2).

26. Wu Y, Jiang T, He X, Shao J, Wu C, Mao W, et al. Global Phylogeography and Genomic Epidemiology of Carbapenem-Resistant bla(OXA-232)-Carrying Klebsiella pneumoniae Sequence Type 15 Lineage. Emerg Infect Dis. 2023;29(11):2246–56.

27. Gao H, Liu Y, Wang R, Wang Q, Jin L, Wang H. The transferability and evolution of NDM-1 and KPC-2 co-producing Klebsiella pneumoniae from clinical settings. EBioMedicine. 2020;51:102599.

28. Hernández-García M, Castillo-Polo JA, Cordero DG, Pérez-Viso B, García-Castillo M, Saez de la Fuente J, et al. Impact of Ceftazidime-Avibactam Treatment in the Emergence of Novel KPC Variants in the ST307-Klebsiella pneumoniae High-Risk Clone and Consequences for Their Routine Detection. J Clin Microbiol. 2022;60(3):e0224521.

29. Peirano G, Chen L, Kreiswirth BN, Pitout JDD. Emerging Antimicrobial-Resistant High-Risk Klebsiella pneumoniae Clones ST307 and ST147. Antimicrob Agents Chemother. 2020;64(10).

30. Jia X, Li C, Chen F, Li X, Jia P, Zhu Y, et al. Genomic epidemiology study of Klebsiella pneumoniae causing bloodstream infections in China. Clin Transl Med. 2021;11(11):e624.

31. Tumbarello M, Raffaelli F, Giannella M, Mantengoli E, Mularoni A, Venditti M, et al. Ceftazidime-Avibactam Use for Klebsiella pneumoniae Carbapenemase-Producing K. pneumoniae Infections: A Retrospective Observational Multicenter Study. Clin Infect Dis. 2021;73(9):1664–76.

32. Shen S, Tang C, Ding L, Han R, Yin D, Yang W, et al. Identification of KPC-112 from an ST15 Klebsiella pneumoniae Strain Conferring Resistance to Ceftazidime-Avibactam. mSphere. 2022;7(6):e0048722.

33. Tatusova T, Ciufo S, Fedorov B, O’Neill K, Tolstoy I. RefSeq microbial genomes database: new representation and annotation strategy. Nucleic Acids Res. 2014;42(Database issue):D553–9.

34. Naas T, Cuzon G, Truong HV, Nordmann P. Role of ISKpn7 and deletions in blaKPC gene expression. Antimicrob Agents Chemother. 2012;56(9):4753–9.

35. Idowu T, Ammeter D, Arthur G, Zhanel GG, Schweizer F. Potentiation of β-lactam antibiotics and β-lactam/β-lactamase inhibitor combinations against MDR and XDR Pseudomonas aeruginosa using non-ribosomal tobramycin-cyclam conjugates. J Antimicrob Chemother. 2019;74(9):2640–8.

36. Jin X, Chen Q, Shen F, Jiang Y, Wu X, Hua X, et al. Resistance evolution of hypervirulent carbapenem-resistant Klebsiella pneumoniae ST11 during treatment with tigecycline and polymyxin. Emerg Microbes Infect. 2021;10(1):1129–36.

37. Yao H, Zhang T, Peng K, Peng J, Liu X, Xia Z, et al. Conjugative plasmids facilitate the transmission of tmexCD2-toprJ2 among carbapenem-resistant Klebsiella pneumoniae. Sci Total Environ. 2024;906:167373.

38. Liu X, Wu Y, Zhu Y, Jia P, Li X, Jia X, et al. Emergence of colistin-resistant hypervirulent Klebsiella pneumoniae (CoR-HvKp) in China. Emerg Microbes Infect. 2022;11(1):648–61.

